# Extracellular release of two peptidases dominates generation of the trypanosome quorum-sensing signal

**DOI:** 10.1101/2021.12.10.472056

**Authors:** Mabel Deladem Tettey, Federico Rojas, Keith R. Matthews

## Abstract

Trypanosomes causing African sleeping sickness use quorum-sensing (QS) to generate transmission-competent stumpy forms in their mammalian hosts. This density-dependent process is signalled by oligopeptides that stimulate the signal transduction pathway leading to stumpy formation. Using mass spectrometry analysis, peptidases released by trypanosomes were identified and, for 12 peptidases, their extracellular delivery was confirmed. Thereafter, the contribution of each peptidase to QS signal production was determined using systematic inducible overexpression *in vivo*, activity being confirmed to operate through the physiological QS signalling pathway. Gene knockout of the QS-active peptidases identified two enzymes, oligopeptidase B and metallocarboxypeptidase I, that significantly reduced QS when ablated individually. Further, a combinatorial gene knockout of both peptidases confirmed their dominance in the generation of the QS signal, with peptidase release of oligopeptidase B mediated via an unconventional protein secretion pathway. This identifies how the QS signal driving trypanosome virulence and transmission is generated in mammalian hosts.

Graphical Abstract

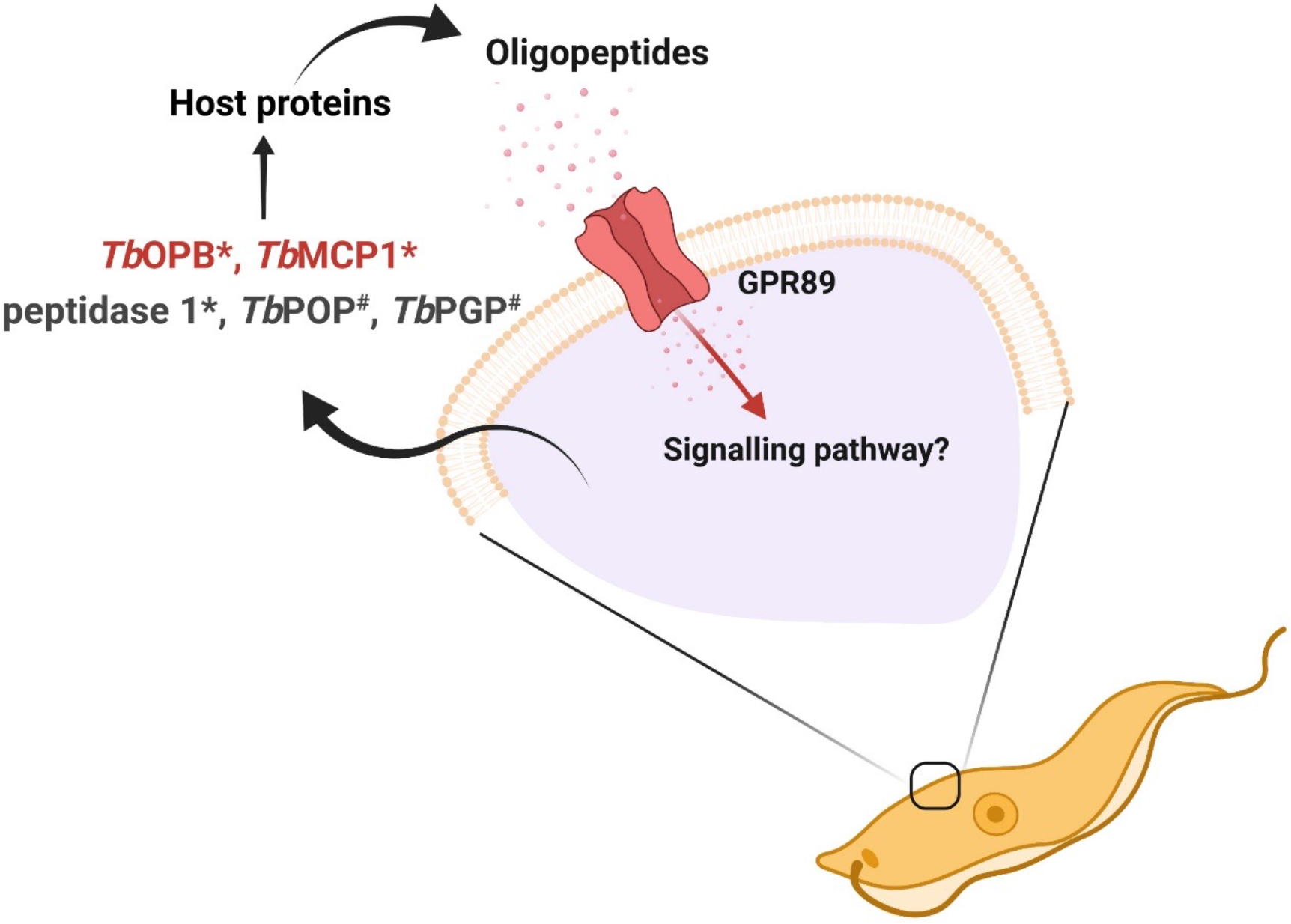

## Introduction

African trypanosomes are responsible for the human disease Human African trypanosomiasis (sleeping sickness) as well as the important livestock disease Animal African trypanosomiasis (nagana)(Buscher et al., 2017; Giordani et al., 2016). These diseases affect health and wealth in afflicted regions of sub-Saharan Africa, infecting thousands of people each year and causing loss to agriculture in the region of $4 billion annually, this acting as a significant driver of poverty. Trypanosomes are transmitted by tsetse flies, with parasite development in the mammalian host contributing to transmission competence(Vickerman, 1985). Specifically, proliferative slender form parasites differentiate to non-proliferative stumpy forms parasites as a preadaptation for tsetse uptake, these stumpy forms exhibiting, for example, mitochondrial activity in preparation for the nutritional environment of the arthropod gut(Dewar et al., 2018; Vickerman, 1965). The development from slender to stumpy forms is a density-dependent phenomenon such that slender forms multiply to sustain the infection whereas arrested stumpy forms dominate at the peak of the infection and throughout the chronic phase of the infection(MacGregor et al., 2011). This, in combination with the parasites’ sophisticated immune evasion strategy, antigenic variation, generates the undulating infection dynamic characteristic of trypanosome infections .

The trypanosome quorum sensing (QS) response is understood in some molecular detail. Firstly, components of the intracellular molecular cascade that transduces the QS signal have been identified by a genome-wide screen selecting for parasites unable to respond to a QS signal mimic after their silencing by RNA interference. This identified several regulators including protein kinases, phosphatases and RNA binding proteins, reflecting the predominant control of trypanosome gene expression at the post transcriptional level(McDonald et al., 2018). At the cell surface, a multimembrane spanning protein TbGPR89 has also been discovered that contributes to QS signalling(Rojas et al., 2018) and this molecule has oligopeptide transport capability. The identification of this molecule highlighted the potential for oligopeptides to act as the external signal driving QS and, indeed, complex oligopeptide mixtures precipitate slender to stumpy differentiation, with synthetic oligopeptides demonstrating specificity of the response. The discovery that short peptides could promote trypanosome differentiation raised the potential that parasite-derive peptidases could generate the extracellular signal for QS. Indeed at least two parasite derived proteases have been identified that are released by bloodstream form parasites and retain activity in the serum of infected animals(Bastos et al., 2010; Morty et al., 2006). These are prolyl oligopeptidase and pyroglutamyl peptidase, and each of these peptidases promotes QS when ectopically over expressed in transgenic bloodstream form parasites(Rojas et al., 2018). In consequence, a model has been developed whereby parasite-derived peptidases generate the QS oligopeptide signal, with their abundance and resultant peptide-generating activity acting as a proxy for parasite density and hence stumpy formation(Rojas and Matthews, 2019). Consistent with this model, basement membrane extracts support efficient parasite differentiation in culture, the enriched extracellular matrix components acting as likely substrates of released proteases (Briggs et al., 2021; Rojas et al., 2021).

A systematic functional screening of trypanosome released proteases has been carried out using targeted RNA interference against each of 30 peptidases encoded in the trypanosome genome(Moss et al., 2015). This revealed that only 1 peptidase produced by the parasites was essential, a signal processing peptidase. Other peptidases were apparently dispensable, and their depletion did not cause growth effects in transgenic parasites grown in culture. However, studies were largely confined to the analysis of parasites grown in culture and the cellular distribution or capacity for release for the targeted peptidases were not analysed, although proteases have been detected in the excretory/secretory products of *Trypanosoma congolense* and *Trypanosoma gambiense*(Bossard et al., 2013). Overall, therefore, the contribution of parasite-derived peptidases to QS is clear, but the enzymes and activities that contribute to driving the process *in vivo* have not been determined.

Here we have analysed the peptidases released by trypanosomes at different stages of development in the mammalian host and during onward differentiation to tsetse midgut forms. Released peptidases have then been evaluated for their ability to contribute to the generation of the QS response, signalled through the identified molecular signalling pathway. In common with an increasing number of parasite-released proteins(Balmer and Faso, 2021), their release occurred through a pathway distinct from classical secretion. In combination, this has identified that two major peptidases dominate the trypanosome QS signalling response, these acting in concert, likely with minor additional contribution from other peptidases, to drive the parasite QS signal generation.

## Results

### Characterisation of the released proteome of bloodstream form trypanosomes

In order to identify proteins released by trypanosome parasites, populations of slender or stumpy parasites were derived from murine infections (stumpy forms) or culture (slender forms; necessary to derive sufficient numbers for analysis), purified and then incubated in serum free Creek’s Medium for 2h. Also, to include proteins released by parasites undergoing differentiation to the next life cycle stage, procyclic forms, stumpy form parasites were incubated in HMI-9 medium containing 6mM cis aconitate and then, after 1 or 4h, parasites were washed and incubated in serum free Creek’s medium containing cis aconitate for 2h (Figure 1A). This provided information on released molecules in bloodstream forms and during the synchronous differentiation of stumpy forms to procyclic forms, after a total of 3h or 6h exposure to the differentiation signal cis-aconitate. After cells were removed by centrifugation, the cell pellet and culture medium derived material was analysed by western blotting with antibody to EF1-alpha to ensure the absence of contamination of the supernatant with cytosolic protein from cell lysis (Supplementary Figure 1A-D). For all samples, three biological replicates were analysed by mass spectrometry, with very good reproducibility between the replicates (slender; >0.947; stumpy >0.769; 3h differentiation >0.942; 6h differentiation >0.890; Figure 1B). We selected proteomic identifications based on protein detection in at least two out of three biological replicates and with at least two distinct peptides identified from the parent protein. Comparing the released proteome of slender and stumpy forms identified 56 proteins common between the two developmental forms, plus 20 that were detected only in slender forms and 29 that were only detected in stumpy forms. Analysis of the differentiating cells by the same criteria revealed 92 that were specifically detected after 3h in cis aconitate, and 52 exclusively detected at 6h, whereas 83 proteins were common between the two differentiation-enriched released proteomes. 19 proteins were common between all of the datasets (i.e., slender, stumpy, 3h and 6h differentiation).

**Figure 1.**
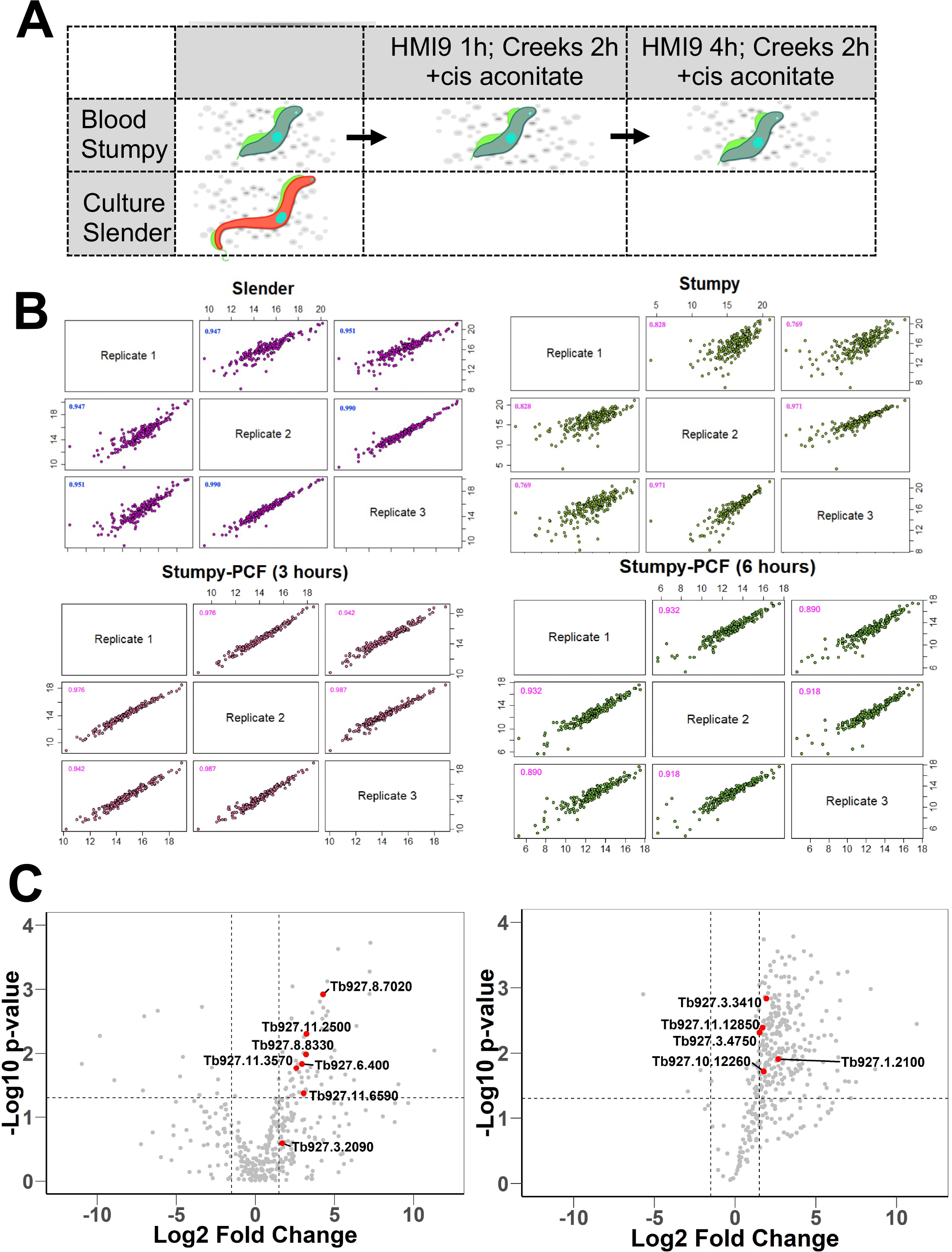
Mass spectrometry analysis of release trypanosome proteins. A. Schematic representation of the samples used for the analysis of the secreted proteome from bloodstream slender forms, stumpy forms or stumpy form parasites undergoing differentiation to procyclic forms. B. Scatter plots showing the reproducibility of biological replicates of secreted material released from slender forms, stumpy forms or cells 3h or 6h into differentiation to procyclic forms. C. Volcano plots of proteins released from either slender or stumpy forms (lefthand panel) or between 3hr and 6hr after the initiation of differentiation between stumpy and procyclic forms (righthand panel). Red dots represent peptidases detected in the dataset; significance thresholds are indicated by dotted lines.

Among the released molecules identified, many peptidases were detected (Supplementary Figure 1E; Figure 1C) and these were of particular interest given their potential contribution to the parasite’s QS response(Rojas et al., 2018) and previous evidence of peptidase release during the course of trypanosome infections(Bastos et al., 2010; Morty et al., 2006). In total 12 peptidases were detected in the released proteome from bloodstream forms and during development to procyclic forms (Supplementary Figure 1E). In a previous global RNA interference screen for protease-related phenotypes(Moss et al., 2015), none of the identified peptidases was detected to have any effect on parasite growth in monomorphic bloodstream form cells in culture. However, monomorphic cells are unable to report on developmental phenotypes in the mammalian host and therefore we specifically explored the released molecules from pleomorphic bloodstream forms further.

### The detected set of 12 peptidases are released by bloodstream form parasites

Initially we sought confirmation that the peptidases detected in the released proteome were indeed derived from intact trypanosomes rather than from damaged or lysed parasites. In the absence of available antibodies to detect each peptidase, we tagged one allele for each peptidase at its endogenous genomic locus, using a Ty1 epitope tag sequence, generating an N-terminal 10 amino acid tag on the encoded protein detectable with the Ty1 specific BB2 antibody(Bastin et al., 1996). The resulting cell lines were assayed for release of the tagged peptidase by subjecting them to the same conditions as used for derivation of the original released proteome, namely incubation in Creek’s medium without serum for 2h, followed by centrifugation to generate the cell pellet and culture supernatant. The resulting material was then analysed for the presence of each tagged peptidase in the pellet or supernatant or for the presence of the abundant cytosolic protein EF1alpha, which was anticipated to remain cell associated if the cells in the assay remained intact. Figure 2A demonstrates that for every peptidase, the tagged protein was distributed between the supernatant and pellet fractions whereas the EF1-alpha signal was exclusively restricted to the pellet, demonstrating cell integrity. This confirmed the peptidases were indeed released from bloodstream form parasites when expressed at physiological levels under their own 3’UTR gene expression control signals. Moreover, the presence of an N-terminal epitope tag suggested a signal sequence was not contributing to their release.

**Figure 2.**
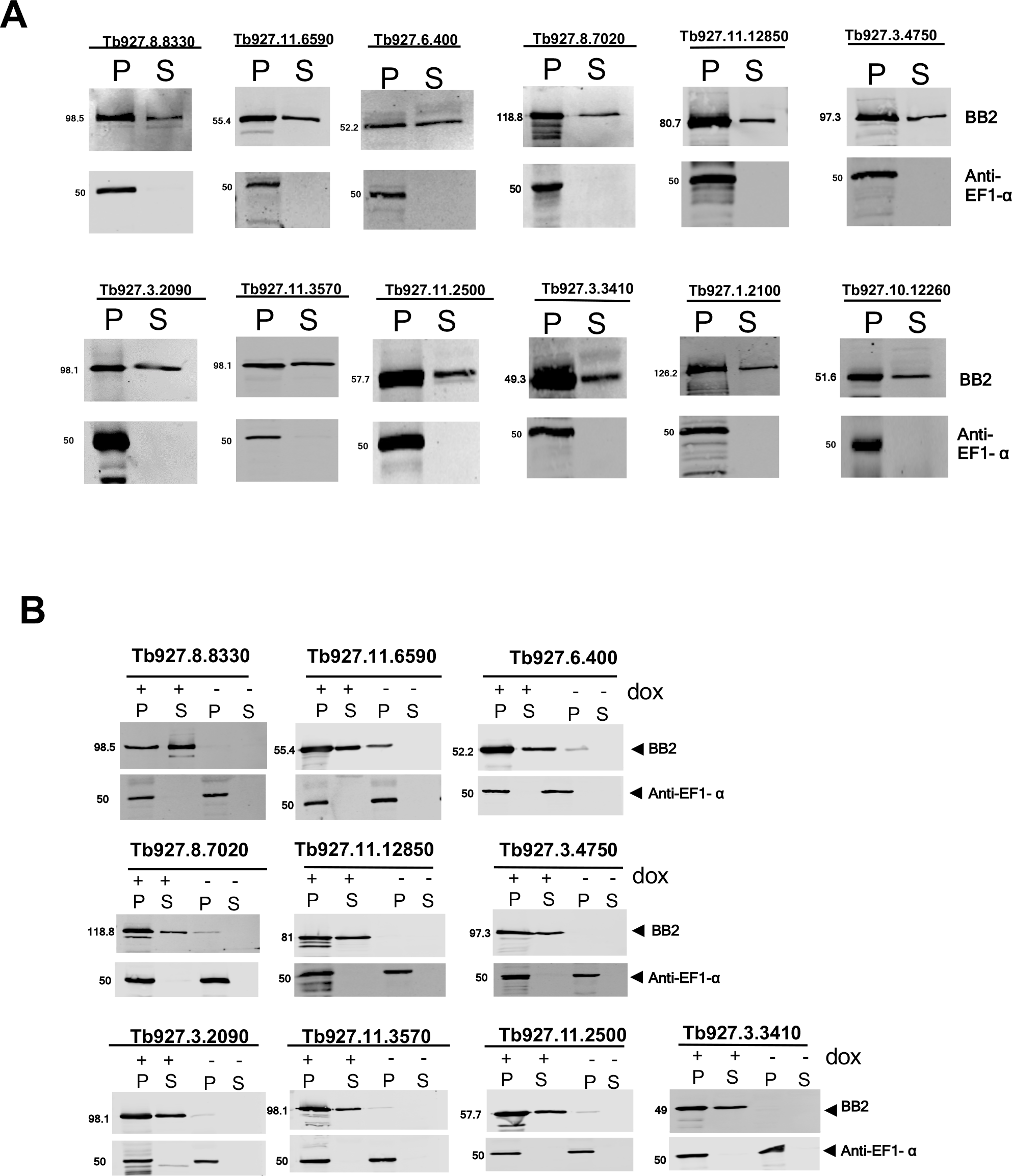
Peptidases identified by mass spectrometry are released from intact trypanosomes. A. Release of identified peptidases by bloodstream form parasites. In each case the identified peptidase was epitope tagged using constructs targeting the endogenous gene locus to ensure close to physiological expression. Resultant cell lines were assayed in serum free Creek’s medium for 2h and then the cell pellet and supernatant isolated by centrifugation. Isolated proteins were monitored for the distribution of each peptidase using the BB2 antibody detecting the Ty1 epitope tag, with the abundant cytosolic protein EF1-alpha used to monitor non-specific escape of cytosolic proteins. B. Inducible ectopic expression of each peptidase identified as released by parasites. In each case N-terminally epitope tagged genes were cloned into a doxycycline inducible expression vector and then the expression and extracellular release of the expressed protein monitored using the epitope tag specific antibody BB2, with the cytosolic protein EF1-alpha used as a control. Released and cell associated proteins were isolated after incubation of parasites in serum free Creek’s medium for 2h and centrifugation.

### Screening released peptides for enhanced development upon ectopic overexpression

To assess whether any of the identified proteases contributed to the generation of the parasite QS signal, each of the released peptidases was engineered for doxycycline inducible ectopic overexpression. Thus, a gene encoding an epitope tagged copy of each peptidase were introduced into a regulatable ectopic expression system in *T. brucei* EATRO 1125 AnTat1.1 90:13, parasites that are pleomorphic *in vivo* and so retain their capacity for QS-driven stumpy formation. Once generated, the cell lines were initially evaluated for the inducible expression of each tagged protein and monitored for their extracellular release. Although cells lines able to inducibly express two peptidases (Tb927.10.12660; Tb927.1.2100) could not be selected, 10 peptidases were successfully expressed, each exhibiting doxycycline inducible expression of the tagged protein (Figure 2B). Furthermore, in each case, the expressed proteins were detected as released from the cells after incubation in serum free Creek’s medium, contrasting with the cytosolic control protein EF1 alpha.

The generated cell lines were then assayed in biological triplicate for phenotypes resulting from their overexpression *in vitro*, with particular focus on any effects on cell growth. Supplementary Figure 2 shows the growth profile of each peptidase-expressing line along with blots confirming the stringent inducible regulation of expression of the tagged proteins. Of the 10 peptidases analysed, ectopic overexpression of two, Tb927.8.8330 and Tb927.11.6590, substantially reduced growth upon induction, with the former resulting in rapid cell death of the parasites (Supplementary Figure 2A). Induction of the expression of Tb927.6.400, Tb927.8.7020 and Tb927.11.12850 reduced the growth of the cells somewhat (Supplementary Figure 2B), whereas Tb927.3.4750 enhanced cell growth when expressed (Supplementary Figure 2C). The remaining four peptidases had minimal effects on the growth of the parasite populations (Supplementary Figure 2D).

Having observed effects *in vitro* we monitored the consequences for parasite differentiation to stumpy forms of ectopic overexpression of each peptidase *in vivo*. Thus, each of the peptidase expressing lines was analysed in triplicate mouse infections, with peptidase expression induced, or not, by provision of doxycycline or 5% sucrose, respectively, to the drinking water of infected animals. Given its strong growth effect *in vitro*, Tb927.8.8330 was not included in the analysis. Infections were monitored over 6 days, with a reduced parasitaemia progression potentially indicative of developmental acceleration in response to the peptidase ectopic overexpression. Figure 3A demonstrates that three peptidases of 9 analysed resulted in substantially premature growth arrest of the parasites *in vivo*, these being Tb927.8.7020 (Peptidase 1), Tb927.11.12850 (Oligopeptidase B) and Tb927.11.2500 (Metallocarboxypeptidase 1). Despite the reduced parasitaemia, the induced parasites were morphologically stumpy and the expression of the stumpy specific marker protein PAD1(Dean et al., 2009) was detectable (not shown). This revealed premature differentiation to stumpy forms was elicited by enhanced expression of each of the three peptidases.

**Figure 3.**
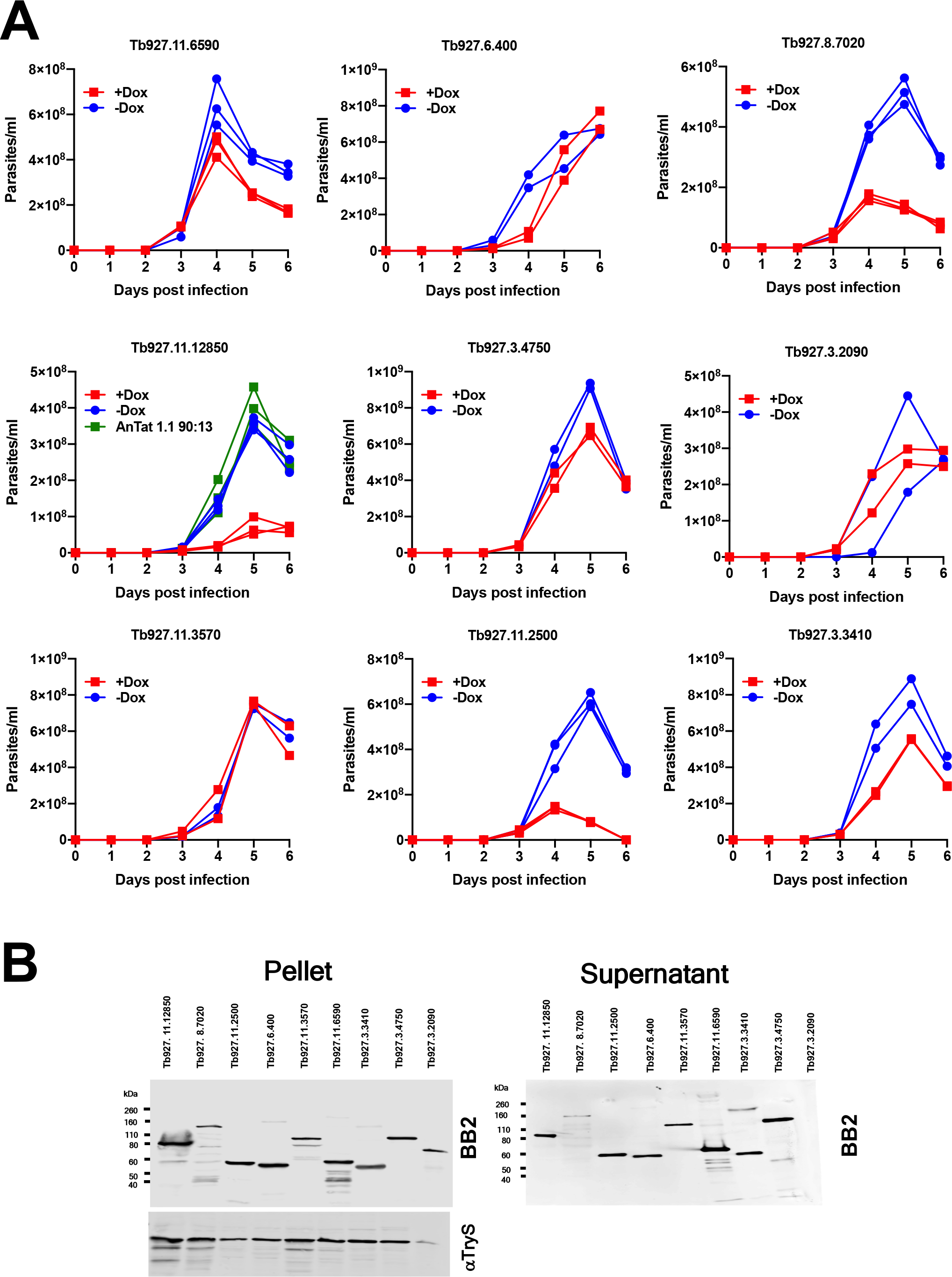
Ectopic expression of three peptidases enhances QS. A. Ectopic overexpression phenotypes of identified peptidases *in vivo*. 1x10^4^ of each parasite line were inoculated into MF1 mice and progression of the parasitaemia monitored after induction of the ectopic peptidase expression from the day of infection. For each peptidase, six mice were infected, 3 being induced and 3 uninduced to express the peptidase via provision of doxycycline in 5% sucrose (induced; red) or 5% sucrose (uninduced; blue) to the drinking water of infected animals. The parasitaemia of the parental *T. brucei* AnTat1.1 90:13 cells was also assayed (green) and is shown in panel Tb927.11.12850 to illustrate its equivalent progression to uninduced parasites. B. Relative expression and extracellular release of each identified peptidase. In each case cell pellet and supernatant were isolated after 2h incubation in Creek’s medium and the epitope tagged peptidase detected using the Ty1-specific BB2 anti body. Loading is indicated by the cytosolic protein Trypanothione synthetase (TyrS). The expression level and extracellular release of the respective peptidases did not correlate with phenotypic effects observed either *in vitro* or *in vivo*.

For all peptidases, the relative level of expression of each, and their relative extracellular release, was compared (Figure 3B). We observed no correlation between those peptidases that induced enhanced differentiation when induced (Figure 3A) and their relative expression or release as assayed by western blotting (Figure 3B). Hence, the premature differentiation that was observed was not a consequence of their expression relative to other peptidases.

### Peptidase expression enhances parasite differentiation in vivo via the QS signalling pathway

We next examined whether the accelerated differentiation caused by expression of the peptidases was stimulated through the characterised quorum signalling pathway. To achieve this, the inducible ectopic expression of each peptidase that promoted accelerated development (Tb927.8.7020, Tb927.11.12850, Tb927.11.2500) was engineered in a cell line deleted for the QS signalling pathway component RBP7(McDonald et al., 2018; Mony et al., 2014). Deletion of this gene renders parasites less able to respond to the QS signal *in vivo* and thereby generate stumpy forms. Figure 4 shows the infection profile of parasites in mice (Figure 4A) with ectopic expression of each peptidase induced or not in the RBP7 null mutant lines. In contrast to cells with an intact QS signalling pathway (Figure 3), in this case the induction of the peptidases did not strongly reduce the parasitaemia *in vivo* and the infections progressed with limited detectable generation of morphologically stumpy forms in either induced or uninduced parasite lines. To ensure that the peptidases were effectively produced and released in the RBP7 null mutant lines, their regulated expression was examined by western blotting, demonstrating inducible expression of each peptidase and its presence in both the cell pellet and supernatant confirming extracellular release (Figure 4B). Despite the high parasitaemias of each infection (exceeding 1x10^9^ parasites/ml), only a low level of PAD1 expression was observed by western blotting (Figure 4B) and immunofluorescence (Figure 4C) and this was not significantly different regardless of the expression of the peptidases.

**Figure 4.**
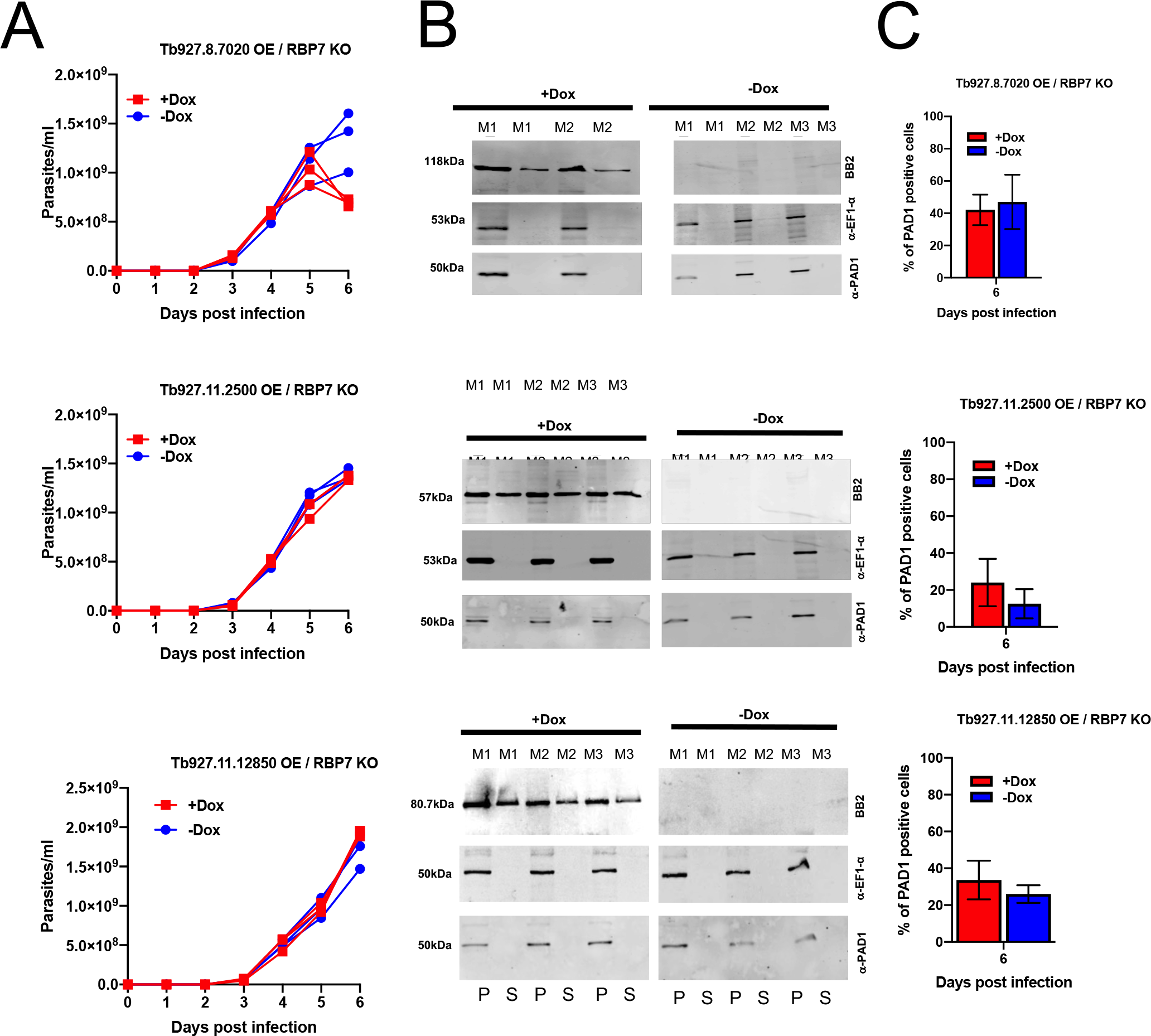
Accelerated differentiation activated by the expression of three peptidase is mediated via the QS signalling pathway. A. Infection profile for mice infected with parasite lines ectopically expressing each peptidase (Tb927.11.2500, Tb927.11.12850 or Tb927.8.7020) under doxycycline regulation with the QS signalling pathway disrupted (⊗RBP7). In each case six mice were infected, with three being provided with doxycycline to induce ectopic peptidase expression (uninduced, blue; induced, red). In all cases the parasitaemia progressed unchecked due to the lack of reception of the QS signal caused by deletion of RBP7. The relative expression of PAD1 scored on a per cell basis by immunofluorescence microscopy is shown for each cell line when harvested on day 6, when parasitaemias exceeded 1x10^9^/ml. B. Confirmation of the inducible expression and extracellular release of each peptidase in an ⊗RBP background. Parasites were harvested from the infections shown in panel A, with three mice being induced for ectopic peptidase expression and three being uninduced. Detection of the stumpy marker PAD1 demonstrated equivalent, low level, stumpy generation in the parasites regardless of the inducible expression of the peptidase. Stringent doxycycline regulated expression is observed in each case. The loading control for the cell pellet was EF1alpha. C. Quantitation of PAD1 expression on day 6 post infection when parastaemias exceeded 1x10^9^/ml. by immunofluorescence assays based on triplicate infection profiles represented in panel A and analysed by western blotting represented in Panel B.

### Peptidase null mutants exhibit delayed differentiation

Having demonstrated that the ectopic overexpression of three peptidases promoted accelerated differentiation of trypanosomes *in vivo*, we explored whether the individual deletion of the genes for each peptidase would reduce differentiation efficiency. Therefore, we exploited CRISPR mediated gene replacement in the *T. brucei* EATRO 1125 AnTat1.1 J1339 cells line which constitutively expresses CAS9(Rojas et al., 2018), to create null mutants for the identified peptidases. Although null mutants for Tb927.8330, Tb927.3.2090 and Tb927.10.12260 were not successfully isolated, null mutants for 8 peptidases were generated. This included the peptidases Tb927.8.7020 (peptidase 1), Tb927.11.2500 (metallocarboxypeptidase 1) and Tb927.11.12850 (Oligopeptidase b), which generated accelerated differentiation when overexpressed. Once gene deletion was confirmed for each of these peptidases (Figure 5A shows the QS-active peptidases), the respective knock out lines were compared to wild type parents in order to assess the parasite growth and level of differentiation to stumpy forms. For all 8 peptidase null mutants, growth *in vitro* was similar to wild type cells (Supplementary figure 3) and for 6 peptidases, including Tb927.8.7020, the progression of the parasitaemia in mice was equivalent to parental cells (Figure 5B and Supplementary figure 4). This indicated their deletion did not affect the fitness or the developmental progression of the parasites *in vivo*. In contrast, null mutants for both Tb927.11.2500 and Tb927.11.12850 both showed a higher total parasitaemia on day 4 post infection compared to parental cells indicating reduced differentiation to stumpy forms *in vivo* (Figure 5B). Nonetheless, although delayed, in both cases the cell lines developed stumpy forms by day 5 of infection, supportive of the signal being generated through the action of more than one peptidase.

**Figure 5.**
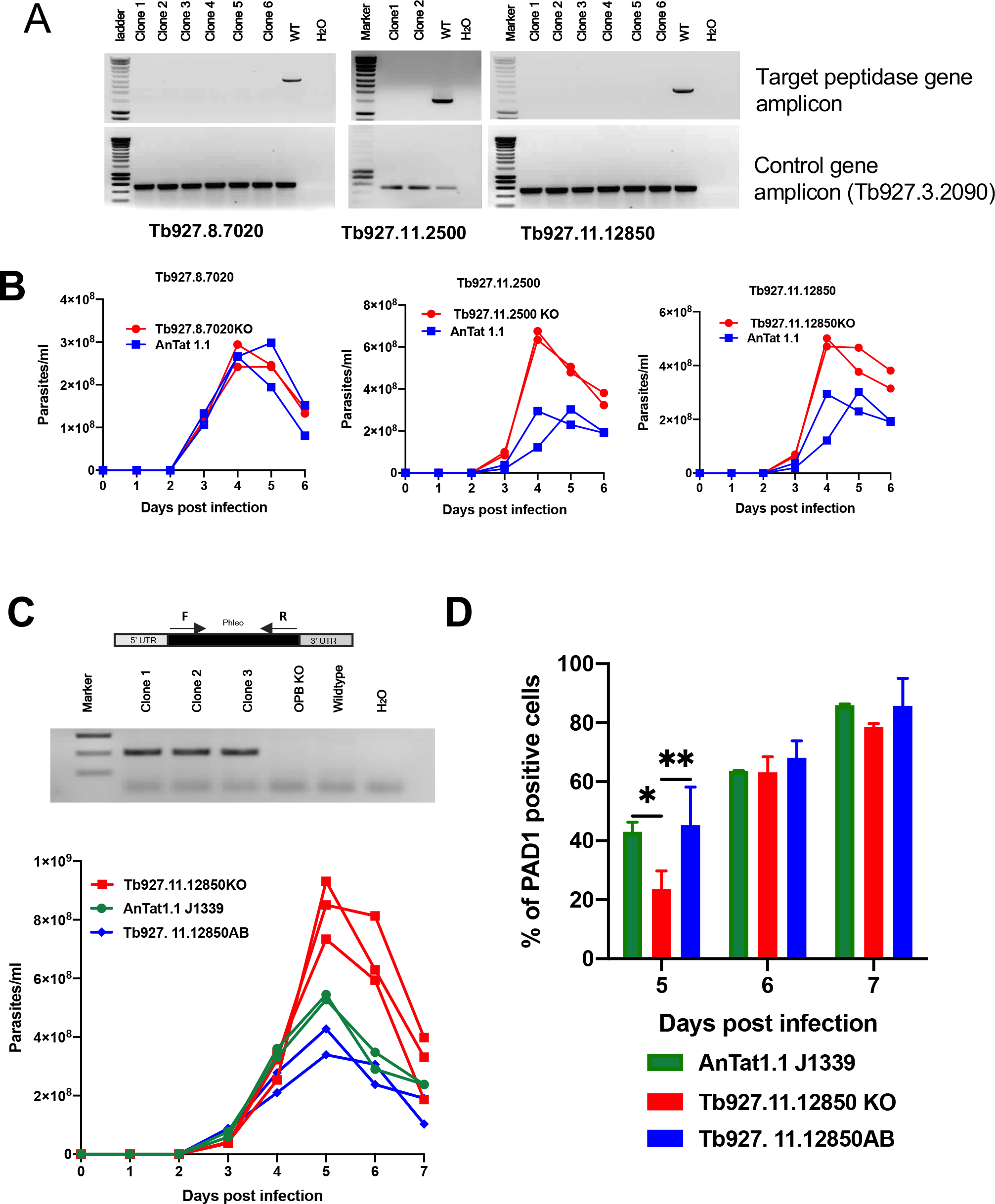
Null mutants for two released peptidases show reduced quorum sensing. A. PCR analysis of genomic DNA derived from null mutants of Tb927.8.7020, Tb927.11.2500 and Tb927.11.12850 analysed using gene specific primers. Several null mutant clones were analysed in each case, together with wild type parental parasites. For each target gene no amplicon was detected with the null mutant clones, whereas a band of the expected size was detected using gDNA from wild type parasites. Integrity of the gDNA in each case was validated using primers targeting a control gene (lower panels). B. Parasitaemias of infections initiated with null mutant line (red) or parental line (blue) for each target gene. Duplicate infections are shown, with Tb927.11.2500 and Tb927.11.122850 showing elevated parasitaemia. Note that the scales differ on the y axis between graphs. C. Upper panel- PCR confirmation of the successful add back of a copy of Tb927.11.12850 to the null mutant. Lower panel- parasitaemias of triplicate infections initiated with parental *T. brucei* AnTat1.1 parasites (green), a Tb927.11.12850 null mutant (red), or the Tb927.11.12850 null mutant containing an add back for the gene (blue). The null mutant exhibited elevated parasitaemia reflecting reduced differentiation, whereas the add back of the Tb927.11.12850 gene restored the growth profile to levels similar to the parental line. D. PAD1 expression of parasites at day 5, 6 and 7 post infections. The null mutant exhibited reduced PAD1 expression at peak parasitaemia compared to the parental cells or add back line. Beyond day 5, PAD1 levels were similar between lines, reflecting that the null mutant exhibited delayed but not abrogated stumpy formation, with stumpy parasites further enriched due to their enhanced tolerance of the developing immune response beyond day 5. Two-way Anova, p<0.05 (*); p<0.01 (**).

To confirm that the reduced differentiation was a consequence of the gene deletion of the peptidases Tb927.1.12850 and Tb927.11.2500, we transfected the null mutant lines to restore expression of each peptidase at its endogenous locus. Once the reintegration of each peptidase gene was validated (Figure 5C shows Tb927.10.12580; Supplementary Figure 5 shows Tb927.11.2500) these ‘add back’ cell lines were assayed for their growth and differentiation *in vivo* in comparison to the parental cell line and each null mutant. Figure 5C and D shows that add back of the peptidase Tb927.11.12850 restored differentiation to levels observed in the parental cell lines, with PAD1 expression being significantly different between the parental and null mutant at peak parasitaemia (day 5; p= 0.014) and between the null mutant and the add back cell line (day 5; p=0.007), but not between the parental and add back lines (Day 5; p=0.929). For Tb927.11.2500, parasites expressing the add back were successfully recovered and the expression of the restored gene confirmed (Supplementary Figure 5A, B) and these cells grew *in vitro* equivalently to wild type cells (Supplementary Figure 5C). However, these parasites were unable to establish infections in mice despite the analysis of several cell lines and in several assays. Consequently, the restoration of stumpy formation could not be unequivocally confirmed in this case.

### Knockout of two peptidases acts combinatorially to reduce QS signalling

Having established that deletion of two peptidases, Tb927.11.12580 (oligopeptidase B) and Tb927.11.2500 (metallocarboxypeptidase 1) reduced differentiation individually we sought to establish if they would operate combinatorially to contribute to the generation of the QS signal. To explore this, a parasite line was created that had deleted both alleles of both peptidase genes (Tb927.11.12580:: Tb927.11.2500) and this line was then compared with the parental line and one of the individual gene knockout lines (Tb927.11.12850 KO) for their growth and differentiation *in vivo*. Figure 6A demonstrated that the deletion of both peptidases generated a parasitaemia of enhanced virulence, exceeding that seen where one peptidase (Tb927.11.12580) alone was absent. Supporting this being a consequence of reduced differentiation efficiency, analysis of the expression of PAD1 revealed that both the single and double peptidase deletion expressed less PAD1 at the peak of the infection on day 5 than the parental line (parent versus Tb927.11.12850 KO, p=0.0004; parent versus Tb927.11.12580 :: Tb927.11.2500, p=0.0002) although the single and double peptidase deletion lines were not significantly different from each other with respect to this marker (p=0.933) (Figure 6B). To confirm that the observed effects on differentiation were not a consequence of multiple transfection rounds of the parasites, Tb927.11.12580 was reintroduced into the double knock out line and the parasitaemia and differentiation examined. Figure 6C demonstrated that re-introduction of the Tb927.11.12580 reduced parasite virulence and enhanced differentiation compared to the double KO mutant, similar to the individual knockout of Tb927.11.12580. On the basis of these data, we conclude that the combined activity of both peptidases, oligopeptidase B and metallocarboxypeptidase 1, contributes to the parasite’s capacity for QS, rather than there being complete redundancy between their action.

**Figure 6.**
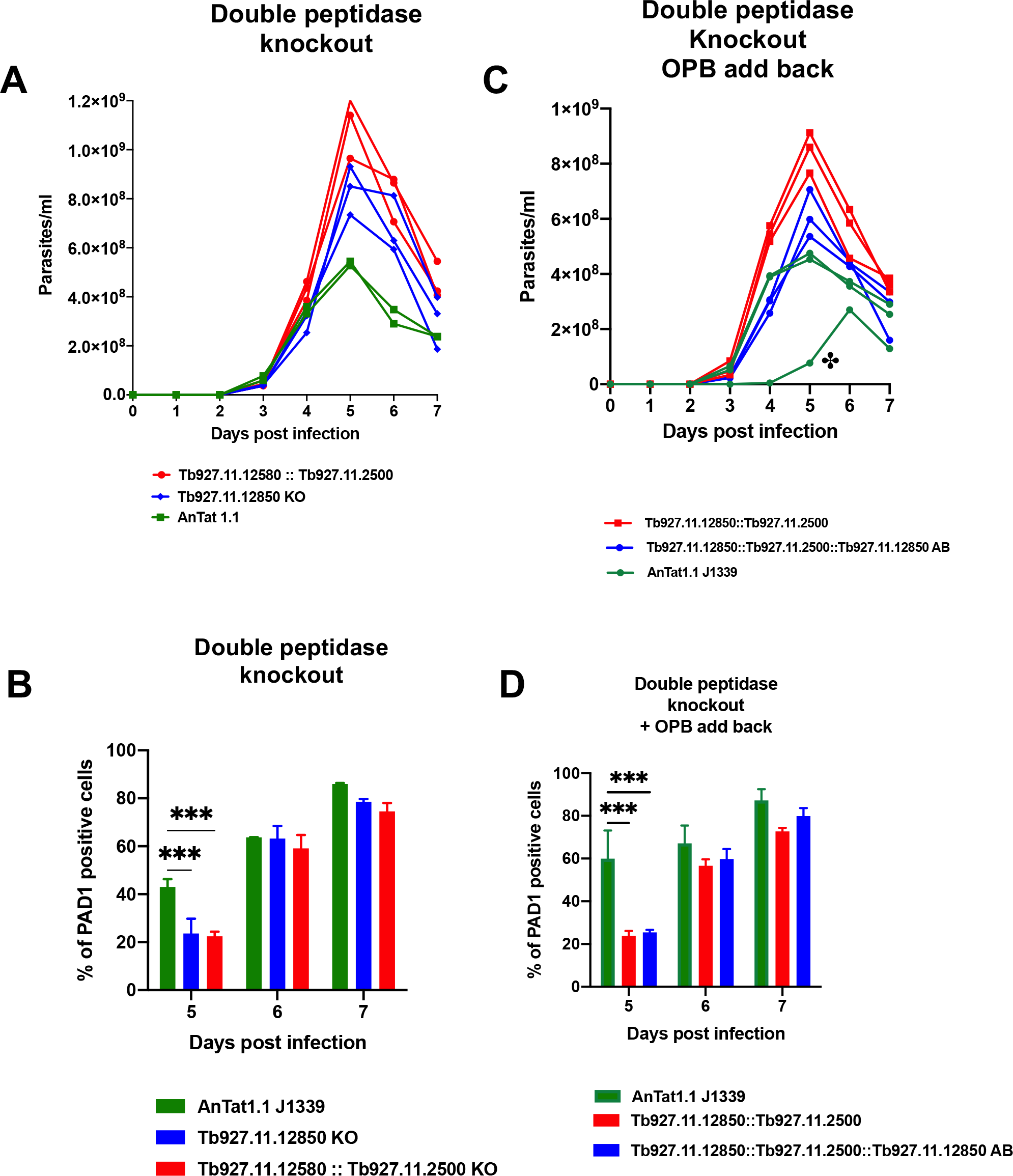
Two released peptidases contribute combinatorially to the generation of the quorum sensing response. A. Infection profile of parental *T. brucei* AnTat 1.1 parasites (green), Tb927.11.12850 null mutants (blue) and Tb927.11.12850 plus Tb927.11.2500 double null mutants (red) *in vivo*. In each case triplicate infections were analysed. The double peptidase KO was most virulent, the single peptidase KO was less virulent but more virulent than the parental line. B. PAD expression of *T. brucei* AnTat 1.1 parasites, Tb927.11.12850 null mutants and Tb927.11.12850 plus Tb927.11.2500 double null mutants on day 5, 6 and 7 of infection. At peak parasitaemia on day 5 both the single peptidase and double peptidase infections inhibited less PAD1 expression than parental cells. Beyond day 5, PAD1 levels were similar between lines, reflecting that the null mutants exhibited delayed but not fully abrogated stumpy formation, with stumpy parasites further enriched due to their enhanced tolerance of the developing immune response beyond day 5. C. Infection profile of parental *T. brucei* AnTat 1.1 parasites (green), Tb927.11.12850 plus Tb927.11.2500 double null mutants (red), and Tb927.11.12850 plus Tb927.11.2500 double null mutants with a restored Tb927.11.12850 gene (blue) *in vivo*. In each case triplicate infections were analysed; one mouse infected with the AnTat1.1 line exhibited abnormally delayed parasitaemia and was discounted from the statistical analysis in panel B. The double peptidase KO was most virulent, whereas the add back less virulent than the double KO but more virulent than the parental line. D. PAD expression of *T. brucei* AnTat 1.1 parasites, Tb927.11.2500::Tb927.11.2500 double null mutants and Tb927.11.2500::Tb927.11.2500 double null mutants with restored Tb927.11.12850 add back on day 5, 6 and 7 of infection. At peak parasitaemia on day 5 both the double peptidase null mutants and the add back mutant infections inhibited less PAD1 expression than parental cells. Beyond day 5, PAD1 levels were similar between lines, reflecting that the null mutants and add back line exhibited delayed but not fully abrogated stumpy formation, with stumpy parasites further enriched due to their enhanced tolerance of the developing immune response beyond day 5. Two- way Anova, p<0.001 (***).

### Peptidase release is mediated via unconventional protein secretion

The predicted protein sequences of the released peptidases characterised in our analyses did not predict the presence of a secretory signal sequence on the N-terminus of the respective proteins. Therefore, we wished to explore whether the QS active peptidases were being released by Golgi-ER mediated classical secretion or by unconventional protein secretion, such as via extracellular vesicles or other uncharacterised pathways. To explore this, the cytological distribution of the peptidases that promoted differentiation were initially assessed via immunofluorescence assay and cell fractionation (Supplementary Figure 6). This revealed that both exhibited a cytosolic location distinct from the ER as visualised by the ER marker BiP, and this was consistent with their enriched fractionation with cytosolic proteins compared with organellar or nuclear protein markers, and previous studies in *T. b. evansi*. To explore the mechanism of release in more detail for oligopeptidase B, cell lines were generated that expressed C-terminally epitope tagged Tb927.11.12850 in cells where a dominant negative mutant of TbSAR1 (TbSAR1-DN), or the wild type protein (TbSAR1-WT) was inducibly expressed, or where Rab11 was inducibly silenced by RNAi. Expression of TbSAR1-DN inactivates the classical secretory pathway by preventing trafficking from the Golgi to the ER whereas Rab11 depletion increases nanotube and EV mediated protein release(Umaer et al., 2018). In contrast to expression of TbSAR1-WT, when TbSAR1 DN was inducibly expressed, cell growth stopped within 12-24h confirming effective expression of the dominant negative protein (Figure 7A), this being consistent with the timescale of previous studies . Analysis of the extracellular release of Tb927.11.12850 during the first 18h of TbSAR1-DN expression however did not reduce the peptidase in the culture supernatant compared to the TbSAR1- WT expressing line, despite cell integrity being sustained on the basis of EF1-alpha cell-association (Figure 7B). Induction of RNAi against TbRab11 also generated a rapid cessation of cell growth (within 24h) (Figure 7C) but here Tb927.11.12850 also continued to be released at levels equivalent to the uninduced cells (Figure 7D).

**Figure 7.**
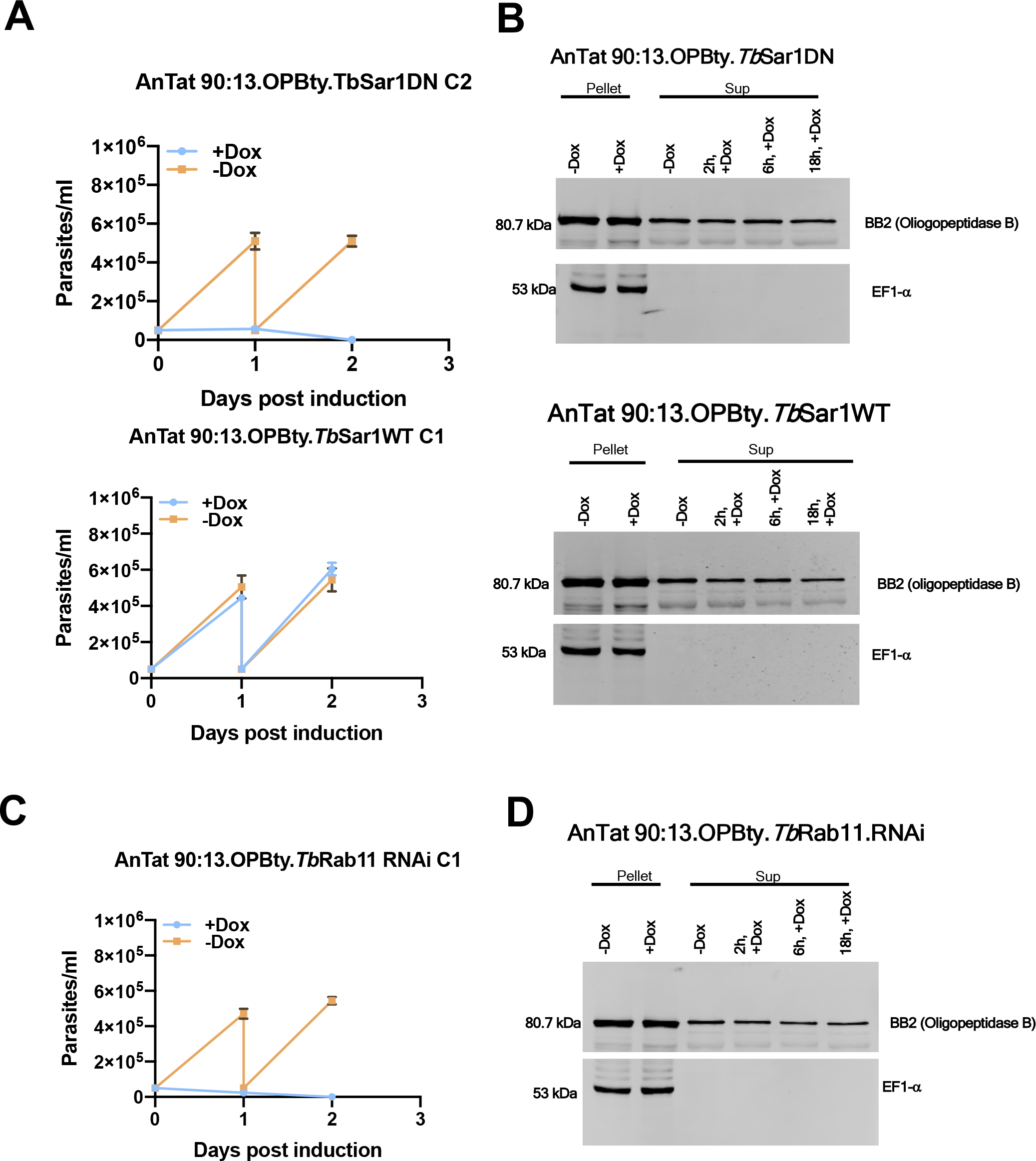
Released peptidases exhibit unconventional protein secretion. A. Growth *in vitro* of *T. brucei* AnTat1.1 90-:13 constitutively expressing epitope tagged Tb927.11.12850 (oligopeptidase B), induced to express dominant negative SAR1 (upper panel) or wild type SAR1 (lowerpanel). Expression of the dominant negative SAR1 induced growth cessation within 24h. B. Extracellular release of epitope tagged Tb927.11.12850 from parasites induced to express dominant negative SAR1 (upper panel) or wild type SAR1 (lower panel). Tb927.11.12850 was released at equivalent levels in each case, whereas the cytosolic control EF1-alpha remained cell associated. C. Growth *in vitro* of *T. brucei* AnTat1.1 90-:13 constitutively expressing epitope tagged Tb927.11.12850, with induced RNAi targeting Rab11. RNAi against Rab11 induced growth cessation within 24h. D. Extracellular release of epitope tagged Tb927.11.12850 from parasites induced to deplete Rab11 by RNAi. Extracellular release of Tb927.11.12850 was sustained, whereas the cytosolic control EF1-alpha remained cell associated.

In combination, this indicated that trypanosome peptidase release was not directed through either the classical secretory pathway or through extracellular vesicle shedding. Instead, an alternative unconventional protein secretion mechanism is demonstrated.

## Discussion

We have explored released peptidases that could contribute to the generation of the trypanosome’s oligopeptide QS signal using a combination of systematic ectopic overexpression and gene deletion. Our analyses identified 12 released peptidases most of which could be deleted without consequence for the parasites *in vitro* or *in vivo*. This is consistent with the RNAi analysis of all trypanosome peptidases by Moss et al.,(Moss et al., 2015) where all but one peptidase was found to be non-essential for the parasite in their systematic screen. However, that analysis used monomorphic parasites that have lost the ability to become stumpy and the study was mostly restricted to phenotypic analysis *in vitro*. By using pleomorphic cell lines able to generate stumpy forms and a combination of ectopic overexpression and gene deletion using CRISPR Cas9 we discovered via *in vivo* analysis that two released peptidases, oligopeptidase B (Tb927.11.12850) and metallocarboxypeptidase 1 (Tb927.11.2500), significantly and individually modulate the levels of parasite QS *in vivo*, suggesting specificity in their QS signal generation. We have previously also shown that prolyl oligopeptidase and pyroglutamyl peptidase can promote parasite differentiation when ectopically overexpressed, but these were not detected as components of the secreted material released by bloodstream from parasites in the current study. Similarly, we show here that Peptidase 1 (Tb927.8.7020) can enhance differentiation when ectopically overexpressed although its deletion does not detectably influence stumpy formation. Hence, several peptidases could contribute to production of the QS signal with considerable redundancy. However, only Oligopeptidase B and metallocarboxypeptidase 1 null mutants affected overall development of the parasites in the bloodstream despite the potential for enhanced prolyl oligopeptidase activity in oligopeptidase B null mutants (Kangethe et al., 2012). Thus, the analysis of individual and combinatorial phenotypes resulting from gene knockout of oligopeptidase B and metallocarboxypeptidase 1 identifies that these molecules are the dominant contributors to the generation of the trypanosome QS signal.

Several analyses of the molecules secreted by African trypanosomes have identified that peptidases are commonly detected in the excretory/secretory material released by parasites in their mammalian host. A comprehensive literature review of these activities released by *Trypanosoma brucei gambiense* provided in Bossard et al., (2013) describes 444 proteins in 12 functional classes, with 10 peptidase families or subfamilies represented. Of the released molecules 134 were bloodstream specific and 10% of those were peptidases. It has been proposed that these molecules contribute to the degradation of peptide hormones contributing to the physiological effects of trypanosome infection. Of those analysed, oligopeptidase B(Morty et al., 2001) and prolyl oligopeptidase have been shown to retain activity in the bloodstream, with oligopeptidase B able to cleave atrial natriuretic factor, vasopressin and neurotensin. In contrast, metallocarboxypeptidase was proposed to modulate the level of vasoactive kinins . The activity of metallocarboxypeptidase 1 was also proposed to act on proteins of the extracellular matrix and it has been suggested their activity against the blood brain barrier could contribute to the parasite’s tropism.

Previous analyses of the release of proteins by African trypanosomes recognised the absence of a signal sequence on many of molecules identified. Classical secretion involves recognition of an N terminal leader sequence and then trafficking from the Golgi through the ER to the flagellar pocket for ultimate release(Link et al., 2021). This process can be disrupted by the expression of dominant negative SAR1 that blocks trafficking between the Golgi and ER, and is rapidly lethal(Sevova and Bangs, 2009). Alternatively, proteins may be released from parasites via nanotube shedding in a process that is enhanced with Rab11 depletion but not signal sequence dependent(Umaer et al., 2018). The release of the oligopeptidase identified in this study did not show evidence of release by either route, suggesting an unconventional protein secretion (UPS) pathway contributes. UPS has been identified as a significant component of extracellular protein release for molecules involved in parasite host interactions, examples including ABC transporter mediated release of acylated parasite proteins (e.g. Leishmania HASP secretion; (Denny et al., 2000)) autophagosome release or other uncharacterised UPS routes(Balmer and Faso, 2021). Supporting a non-ER based release of the QS- peptidases identified here, both peptidases exhibited cytosolic distribution by immunofluorescence microscopy and cell fractionation.

Trypanosomes undergo stumpy formation in the bloodstream but also adipose tissue and skin, and in each compartment, they can comprise the dominant developmental form in chronic infections (Capewell et al., 2016; MacGregor et al., 2011; Trindade et al., 2016). In the bloodstream, stumpy formation is related to parasite density due to the enhanced QS signal generation provided by large numbers of parasites releasing peptidases and so generating the oligopeptide signal driving the developmental response(Matthews, 2021). In livestock infections, however, parasitaemias are lower and numbers may not drive the QS response in the bloodstream as effectively as in high parasitaemia experimental infections in mice. However, for parasites constrained in the tissues, absolute parasite numbers are less relevant for QS than the local concentration of parasites and their released peptidases, which can generate oligopeptide signals proximally to the compartmentalised parasites(Rojas et al., 2018). In this low flow environment, QS can generate stumpy forms at low parasitaemia and as a high proportion of the local parasite concentration. In this scenario, stumpy forms dominate favouring parasite transmission, resolving the so-called ‘transmission paradox’ where parasite spread is maintained despite low overall parasite numbers in the host as determined by bloodstream parasitaemias(Capewell et al., 2019). These local stumpy populations, observed in skin and adipose tissue, have the ability to sustain disease spread because small numbers of parasites can infect tsetse flies, at least under laboratory conditions using permissive flies(Matthews and Larcombe, 2021; Schuster et al., 2021). Our discovery that oligopeptide signals from the action of oligopeptidase B and metallocarboxypeptidase 1 in infections provides a coherent picture of how parasites constrained in the tissues can promote the formation of transmissible stumpy forms, including within a tissue environment able to support disease spread within the physiological context of parasites in their natural host setting.

## Materials and Methods

### Trypanosome cell lines

*T. brucei* EATRO (East African Trypanosomiasis Research Organisation in Tororo, Uganda) 1125 AnTat 1.1 90:13 cells and AnTat 1.1 J1339, both pleomorphic cell lines, were used in this study. Pleomorphic cell lines are able to differentiate to the various life cycle forms of the parasite.

The pleomorphic bloodstream forms of the parasite were cultured in HMI-9 medium supplemented with 20% v/v Fetal Bovine Serum (FBS) at 37°C with 5% CO_2_. They were maintained between 1x10^5^ and 1x10^6^ cells/ml and passaged every 24 to 48h in a vented culture flask to allow the diffusion of CO_2_ into the flask.

### Mouse infections

MFI mice were used in this study and inoculated with various *T. brucei* cell lines intraperitoneally. The parasitaemia was monitored daily from day three post-infection. The appropriate life cycle forms of the parasite were harvested by collecting blood from trypanosome-infected mice by cardiac-puncture. The parasites were then purified by separation on a diethyl aminoethyl (DEAE) cellulose anion exchange column . The parasites were subsequently counted using the Neubauer haemocytometer and then washed with Phosphate Buffered Saline-Glucose (PSG) (44mM NaCl; 57mM Na_2_HPO_4_3mM KH_2_PO_4_; 55mM glucose, pH 7.8) by centrifugation at 1,600 x g for 10 minutes. The cells were resuspended in an appropriate volume of pre-warmed HMI-9 media.

All work was carried out under a UK home office licence (P262AE604) that had been approved after local ethical review.

### Detection of released extracellular proteins

To isolate released parasite-derived material from cultured bloodstream forms, 2x10^7^ trypanosomes were washed with PSG at 1,600 x g for 5 minutes twice and incubated in Creek’s Minimal Medium (CMM) without serum for 2h at 37°C, 5% CO_2_ . After the 2-hour incubation, the cells were centrifuged at 1,054 x g for 10 minutes and the supernatant removed. This was spun again at 1,054 x g for 10 minutes to ensure the removal of all parasites from the supernatant. An aliquot of 100 μl of the supernatant was analysed for trypanosome EF1-alpha (EF1〈) using a western blot. The remaining protein was concentrated using cold acetone precipitation method. Briefly, the supernatant was mixed with pre-cold acetone 4x the volume of the supernatant and incubated at -20°C overnight. This was spun the following day at 15,000 x g for 10 minutes and the pellet dried at room temperature for 30 minutes to allow the acetone to evaporate. The precipitated protein samples were then submitted to the Centre for Synthetic and Systems Biology, the University of Edinburgh for MS/MS analysis. Triplicate samples were prepared.

To isolate stumpy derived material, the stumpy forms of *T. brucei* EATRO 1125 AnTat 1.1 90:13 were harvested from mouse infections and purified by DE52 anion exchange chromatography. The parasites were immediately washed with PSG and incubated in HMI-9 medium supplemented with 20% v/v FBS for 2h at 37°C, 5% CO_2_ to allow the parasites to adapt to the *in vitro* condition. After the adaptation period, 2x10^7^ cells were washed by centrifugation with PSG and incubated in CMM without serum for 2h at 37°C, 5% CO_2_ to allow for secretion/release of proteins.

For material derived from parasites undergoing differentiation to procyclic forms, *T. brucei* EATRO 1125 AnTat 1.1 90:13 stumpy forms harvested and purified from infected mice were washed with PSG and incubated in HMI-9 medium supplemented with 20% v/v FBS and 6mM of cis-aconitate to induce differentiation to the procyclic forms for one and 4h at 37°C, 5% CO_2_. After each incubation period, 2x10^7^ cells were washed by centrifugation with PSG and incubated in CMM plus 6mM cis-aconitate without serum for 2h at 37°C, 5% CO_2_. The supernatant was collected after the incubation period and processed the same way as for slender and stumpy forms.

### Mass spectrometry

Samples were run on a pre-cast Bolt™ 4-12% Bis-Tris Plus gel (Invitrogen) for 5 minutes before overnight in-gel trypsin digestion. Peptide extracts were dried by Speedvac and the dried peptide samples were re-suspended in MS-loading buffer (0.05% trifluoroacetic acid in water) then filtered using a Millex filter before HPLC-MS analysis. The analysis was performed using an online system of a nano-HPLC (Dionex Ultimate 3000 RSLC, Thermo-Fisher) coupled to a QExactive mass spectrometer (Thermo-Fisher) with a 300 µm x 5mm pre-column (Acclaim Pepmap, 5 µm particle size) joined with a 75 µm x 50 cm column (Acclaim Pepmap, 3 µm particle size). Peptides were separated using a multi-step gradient of 2–98% buffer B (80% acetonitrile and 0.1% formic acid) at a flow rate of 300 nl/min over 90 minutes. Raw data from MS/MS spectra were searched against a *T. brucei* database using MASCOT (version 2.4). The parameters used in each search were: (i) missed cut = 2, (ii) fixed cysteine carbamidomethylation modification, (iii) variable methionine oxidation modification, (iv) peptide mass tolerance of 10 ppm, (v) fragment mass tolerance of 0.05 Da. Search results were exported using a significance threshold (p-value) of less than 0.05 and a peptide score cut off of 20.

### Immunofluorescence

Cells were either fixed as blood/culture smears air dried on microscope slides and then immersed with ice cold methanol, or were paraformaldehyde fixed in suspension. Specifically, 2x10^6^ cells were pelleted at 2450 x g for 5 minutes and washed once with cold PBS. The cells were then fixed by resuspending them in in 125 μl of cold PBS and 125 μl of 8% paraformaldehyde for 10 minutes on ice. Thereafter, fixed cells were resuspended in 130 μl of 0.1 M glycine in PBS and incubated at 4**°**C at least overnight. After this incubation period, the cells were spun, and the pellet resuspended at 1x10^5^ cells per 10 μl of 1x PBS. Cells were adhered to Polysine**®** slides (VWR, 631-0107) in areas demarcated using an ImmEdge hydrophobic barrier pen (Vector laboratories, H-4000) for 1 hour at room temperature in a humidity chamber. After the 1 hour incubation, the excess PBS was removed from the wells using an aspirator. For the identification of proteins other than surface proteins, cells were permeabilised with 20 μl of 0.1% triton in PBS for 2 minutes. The wells were washed with PBS and blocked with 2% BSA for 1 hour at room temperature. The slides were then washed in 1x PBS for 2 minutes and incubated with 50 μ of appropriate primary antibody (diluted in 2% BSA/PBS, 1:5 for BB2 antibody, 1:1000 for PAD1 antibody) for 1 hour at 37**°**C or 4°C overnight in a humidity chamber. The slides were washed five times, 5 minutes each with 1x PBS followed by 50 μl of appropriate secondary antibody incubation (diluted in 2% BSA/PBS, α-mouse Alexa fluor 568 1:500, α-rabbit Alexa fluor 488 1:500). After the secondary antibody incubation, the slides were washed five times, 5 minutes each and stained with 10 μg/ml 4, 6-diamidino2-phenylindole (DAPI, Life Technologies) for 2 minutes, then washed with 1x PBS for 5 minutes and then mounted with 50 μl Mowiol containing 2.5% 1,4-diazabicyclo [2.2.2] octane (DABCO). The slides were covered with coverslips after the Mowiol addition and analysed on a Zeiss Axioskop 2 plus or Zeiss Axio Imager Z2. QCapture Suite Plus Software (version 3.1.3.10, https://www.qimaging.com) was used for image capture and ImageJ for image analysis.

### Western blotting

Cells were pelleted by centrifugation at 1,646 x g for 5 minutes and then washed with PSG. The resulting cell pellets were resuspended in l00 μl of 1 mM ice-cold Tosyl-L-lysyl-chloromethane hydrochloride (TLCK) and then incubated on ice for 5 minutes and then at 37°C for 15 minutes. After which appropriate volume of Laemmli buffer (62.5 mM Tris-HCl pH6.8, 2% SDS, 10% glycerol, 0.002% Bromophenol blue, 5% β- mercaptoethanol) was added. The samples were either boiled at 95°C for 5 minutes to denature the proteins before storage at -20°C or they were directly stored at -20°C until used.

Proteins resolved by SDS PAGE were transferred to a nitrocellulose transfer membrane (GE Healthcare Amersham™ Protran™) at 80 V for 75 minutes or 15 V overnight at 4°C using a Bio-Rad wet transfer apparatus. The membranes were then stained with ponceau for about five mins to ensure the proteins were properly transferred and then washed with distilled H_2_O several times. The membranes were blocked with LI-COR Odyssey blocking buffer for one hour at room temperature or 4°C overnight and then probed with the appropriate primary antibody for one hour 30 minutes at room temperature or 4°C overnight. The membranes were washed three times, 10 minutes each in Tris-buffered saline with 0.05% Tween (TBST) and then incubated in an appropriate secondary antibody for one hour at room temperature, diluted in 50% LI-COR Odyssey blocking buffer and 50% TBST (1:5000). The membranes were further washed three times, 10 minutes each in TBST and the proteins visualized by scanning the blots using a LI-COR Odyssey Imager, which uses an infrared laser to detect the fluorochrome on the secondary antibody.

### Gene manipulation using CRISPR-Cas9

Gene-specific primers for the amplification of the pPOT plasmid and the small guide RNA (sgRNA) scaffold (G00) were designed using the program at (www.leishgedit.net/Home.html). 0.5 μM of gene-specific reverse and forward primers, 0.2 mM of dNTPs, PHIRE II polymerase were mixed in 1x PHIRE II reaction buffer, the total volume of 150 μl. PCR cycling conditions were 98°C for 30 seconds followed by 98°C for 10 seconds, 59°C for 30 seconds, 72°C for 15 seconds for 35 cycles and then a final extension at 72°C for 5 minutes. Then precipitated material was then used for parasite transfection.

Thus, 3 x 10^7^ cells were used for each transfection. For CRISPR/Cas9 technology, cell lines constitutively expressing cas9 enzymes were used . Cells were centrifuged at 1,600xg for 5 mins and washed by centrifugation with 1 ml of 1x Tb-BSF buffer (90mM NaCl, 5mM KCl, 0.15mM CaCl, 50mM HEPES, pH 7). The cells were then resuspended in 150 μl of 1x Tb-BSF buffer, mixed with 10 μg of precipitated DNA and then transferred into a transfection cuvette. The cells were then electroporated using the Amaxa Nucleofector II and programme Z-001. Cells were recovered in 25 ml of HMI-9 supplemented with 20% FBS v/v for at least 5h, after which they were selected with the appropriate drugs (Blasticidin, 2.5 µg/ml; Phleomycin 0.5 µg/ml).

### Statistical analyses

Graphical and statistical analyses were carried out in GraphPad Prism version 9 (GraphPad Software, La Jolla, California, USA, https://www.graphpad.com) by two- way repeated-measures ANOVA test followed by Tukey or Bonferroni post hoc analysis. For individual experiments, n values are included in the Figure legend; graphs provide mean values ± SD. P values of less than 0.05 were considered statistically significant.

## Acknowledgements

Mass spectrometry was carried out on equipment awarded to the Centre for Immunity, Infection and Evolution (Wellcome Trust award 095831/Z/11/Z). The proteomics analyses were carried out by the EdinOmics research facility at the University of Edinburgh and we particularly acknowledge the assistance of Lisa Imrie. We also thank Dr Nisha Philip (University of Edinburgh) and Professor Jay Bangs (Buffalo University, New York) for invaluable comments and input. Work in Keith Matthews laboratory is funded by the Wellcome Trust (103740/Z/14/Z, 221717/Z/20/Z). Mabel Deladem Tettey was supported by the Darwin Trust of Edinburgh. This research was funded in whole, or in part, by the Wellcome Trust [103740/Z/14/Z, 221717/Z/20/Z]. For the purpose of open access, the author has applied a CC BY public copyright licence to any Author Accepted Manuscript version arising from this submission.

## Author contributions

**MT:** Conceptualisation, conducted the experiments, analysed the data, wrote the manuscript

**FR:** Conceptualisation, conducted the experiments, analysed the data, wrote the manuscript, provided supervision

**KM** Conceptualisation, analysed the data, wrote the manuscript, provided supervision, project administration, funding acquisition

---

**Supplementary Figure 1.**
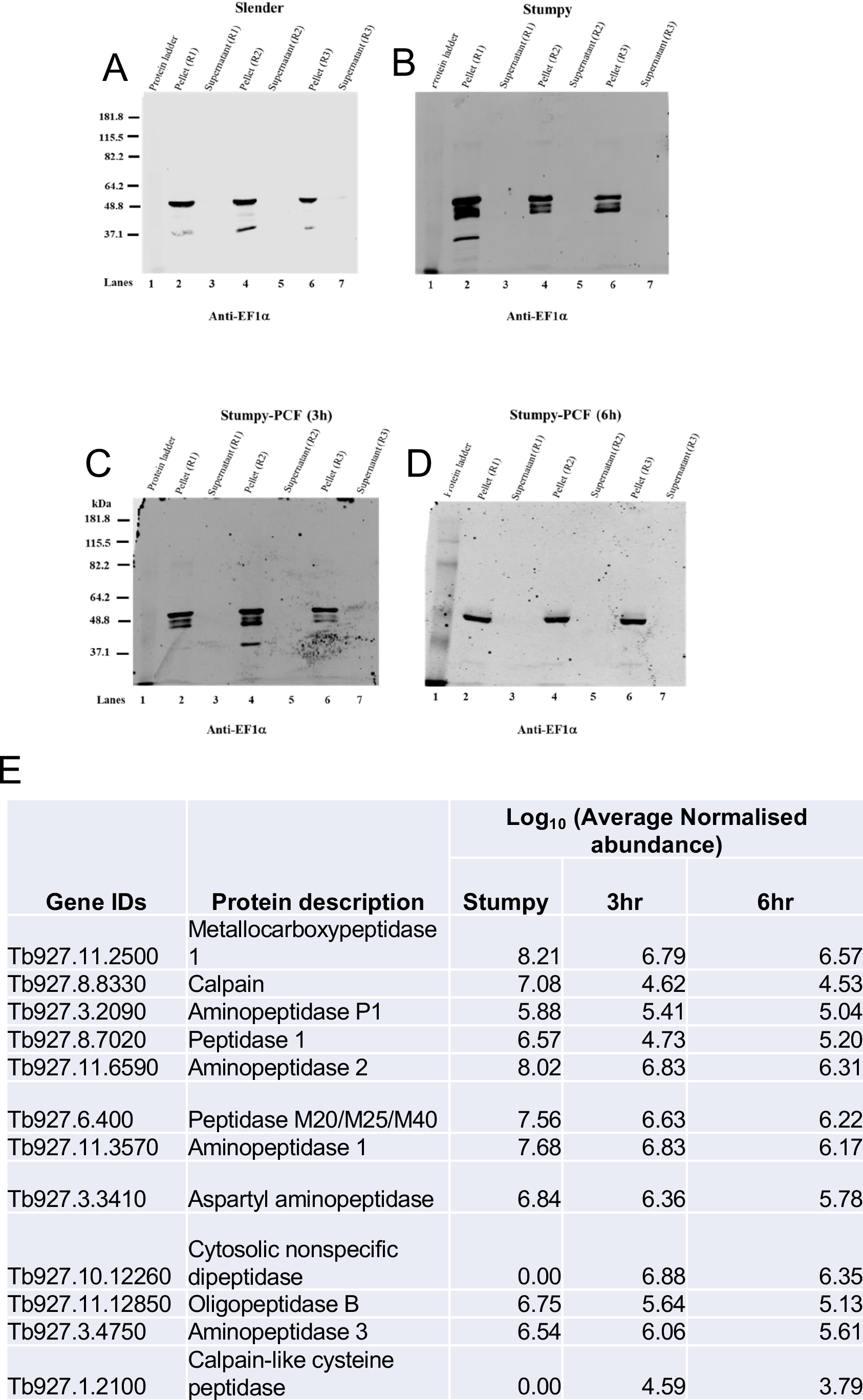
Analysis of trypanosome-released proteins identified many peptidases. A. Slender supernatant, B. stumpy supernatant, C. Stumpy-procyclic differentiating cells supernatant (1h in HMI-9, 2h in CMM), D. Stumpy-procyclic differentiating cells supernatant (4h in HMI-9, 2h in CMM). Three biological replicates are shown in each case, with EF1-alpha signal being detected in the cell pellet but not cell supernatant fractions for each, reflecting the cell integrity of the material used to analyse the secreted/release material from each parasite sample. These samples were used for mass spectometry analysis. Panel E shows peptidases identified in the study.

**Supplementary Figure 2.**
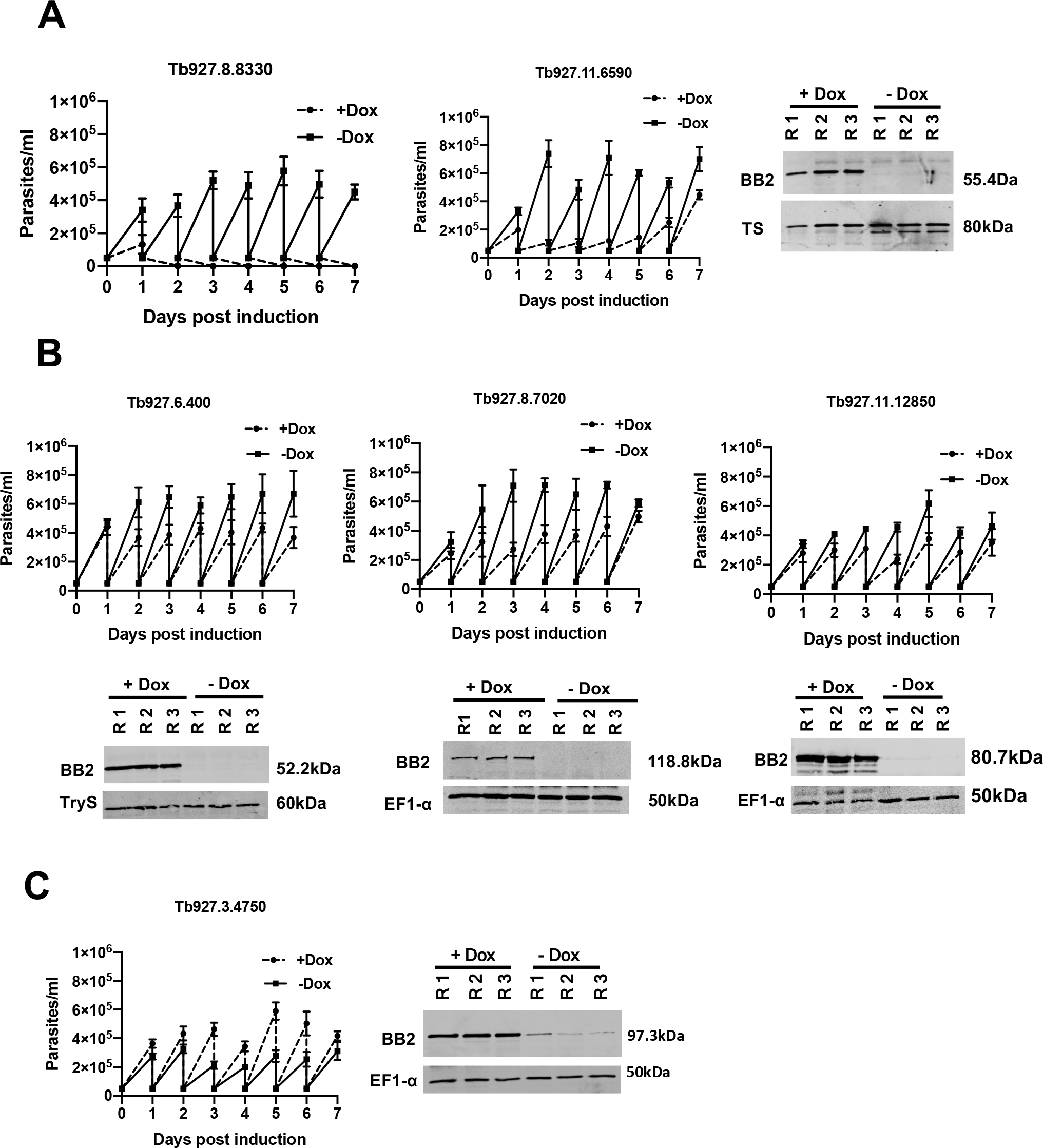
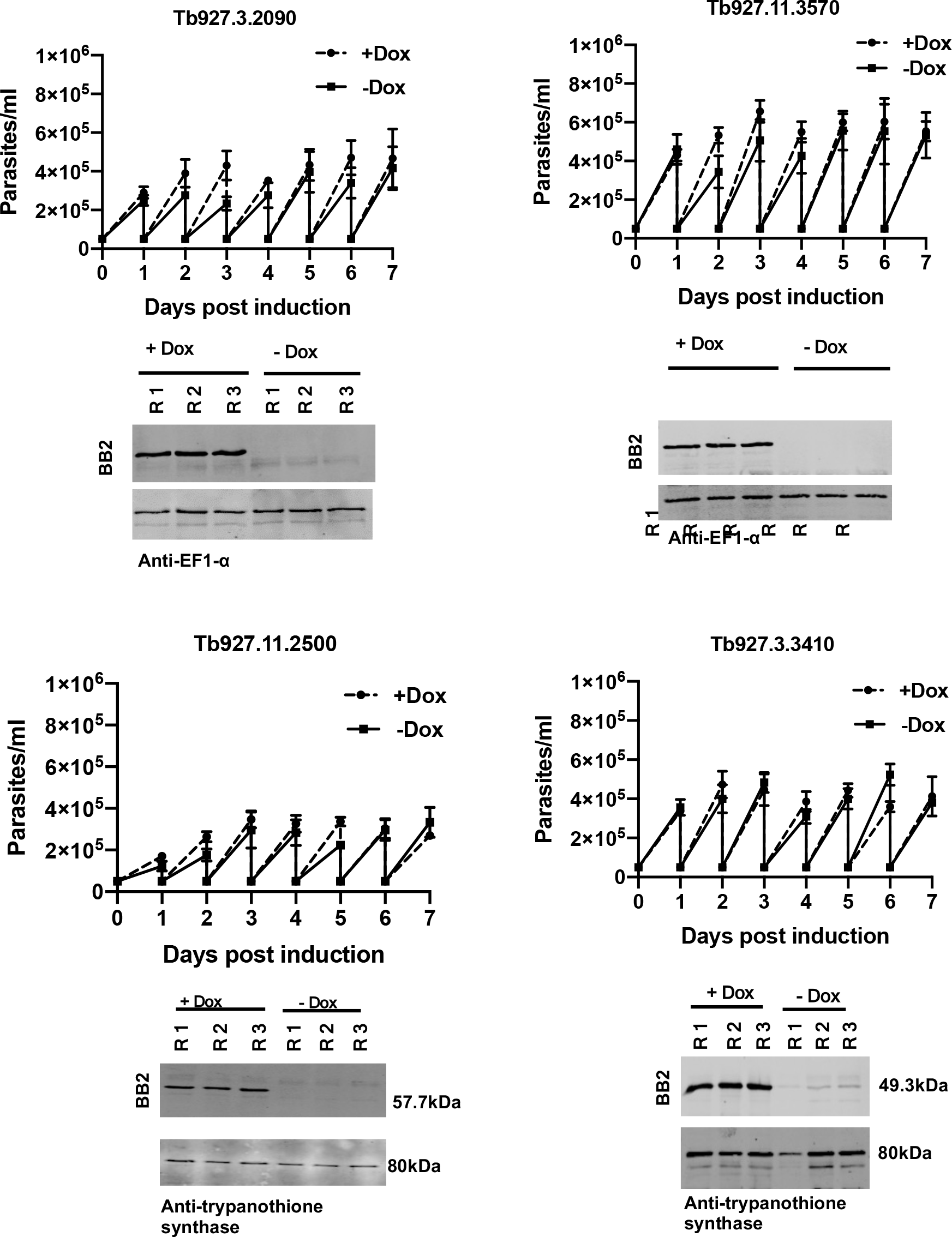
*In vitro* growth profile of parasite cell lines induced, or not, to ectopically overexpress each peptidase. Growth profile of parasites induced (dotted lines), or not (solid lines), to ectopically overexpress each peptidase. On each day, parasites were diluted to 1x10^5^ parasites per ml; each growth profile represents the mean and standard deviation of three biological replicates. The accompanying panels demonstrate the inducible expression of each peptidase detected using the Ty1-eptitope tag specific BB2 antibody. Loading is shown by the abundance of EF1-alpha in each sample. No accompanying western blot is provided for Tb927.8.8330 because the rapid cell death observed after expression prevented the isolation of cellular proteins. A. Peptidases whose expression strongly limits growth of the parasites B. Peptidases whose expression reduces growth of the parasites moderately C. Peptidase whose expression enhances growth of the parasites D. Peptidases that have little discernible effect on the growth of the parasites when ectopically expressed.

**Supplementary Figure 3.**
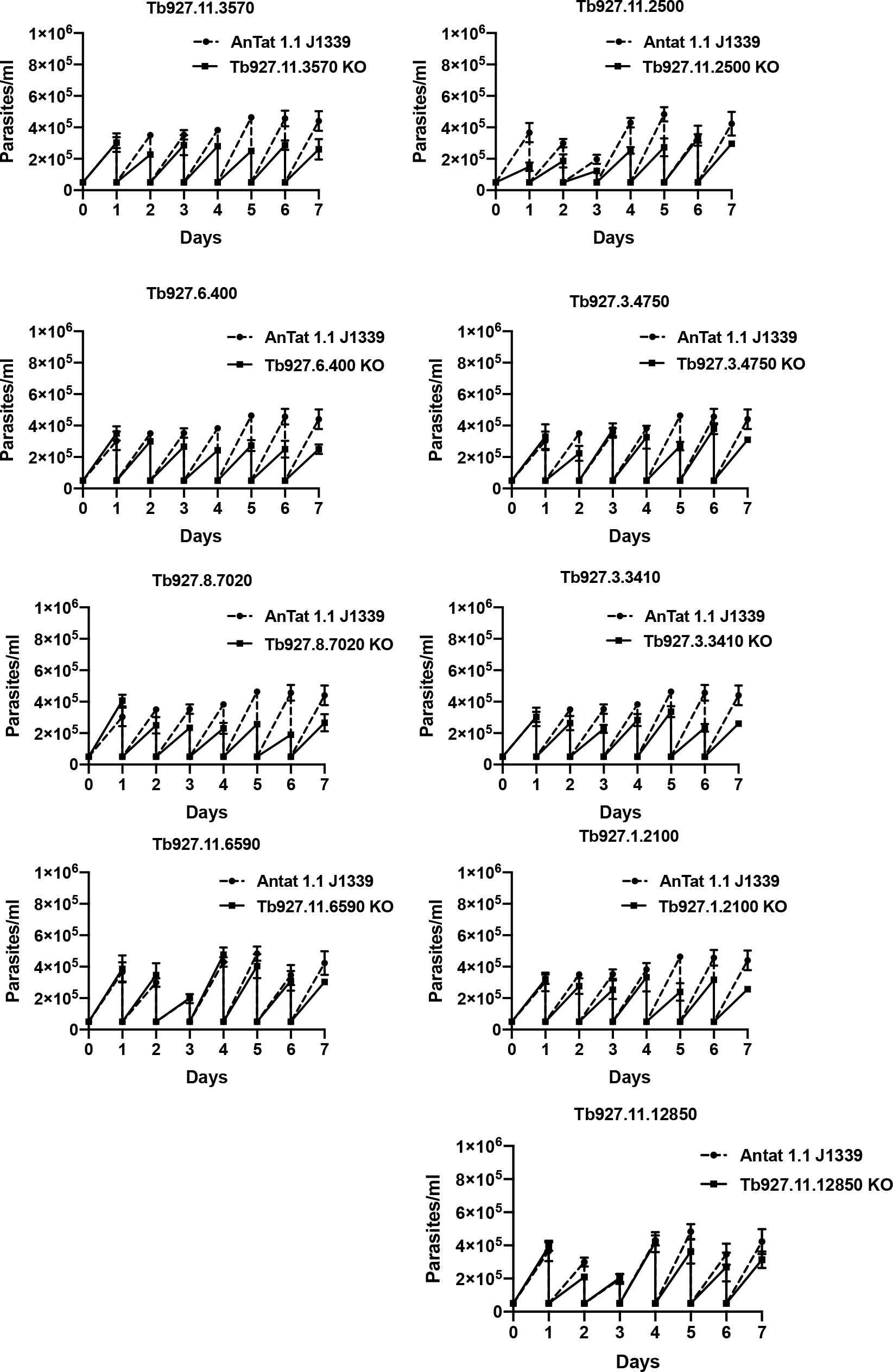
*In vitro* growth of null mutant parasite lines for each identified peptidase. *In vitro* growth profile of each peptidase null mutant line versus growth of parental cell line (*T. brucei* AnTat1.1 J1339). Each day parasites were diluted to 1x10^5^ parasites/ml. Each growth profile represents the mean and standard deviation of three biological replicates.

**Supplementary Figure 4.**
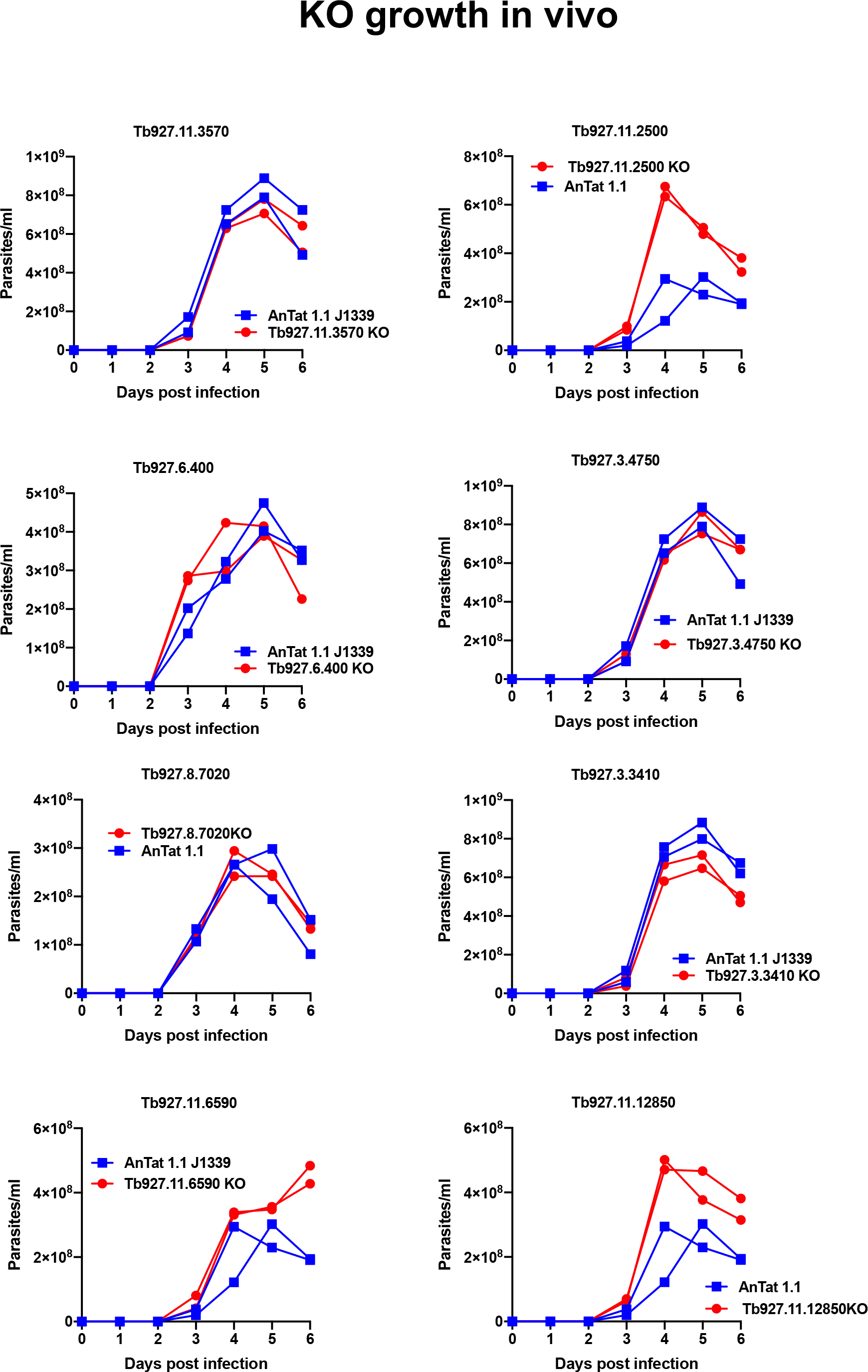
*In vivo* growth of null mutant parasite lines for each identified peptidase. *In vivo* growth profile of each peptidase null mutant line (red lines) versus growth of the parental cell line (*T. brucei* AnTat1.1 J1339) (blue lines). In each case two mice were infected with the null mutant line and two with the parental line and infections carried out in parallel.

**Supplementary Figure 5.**
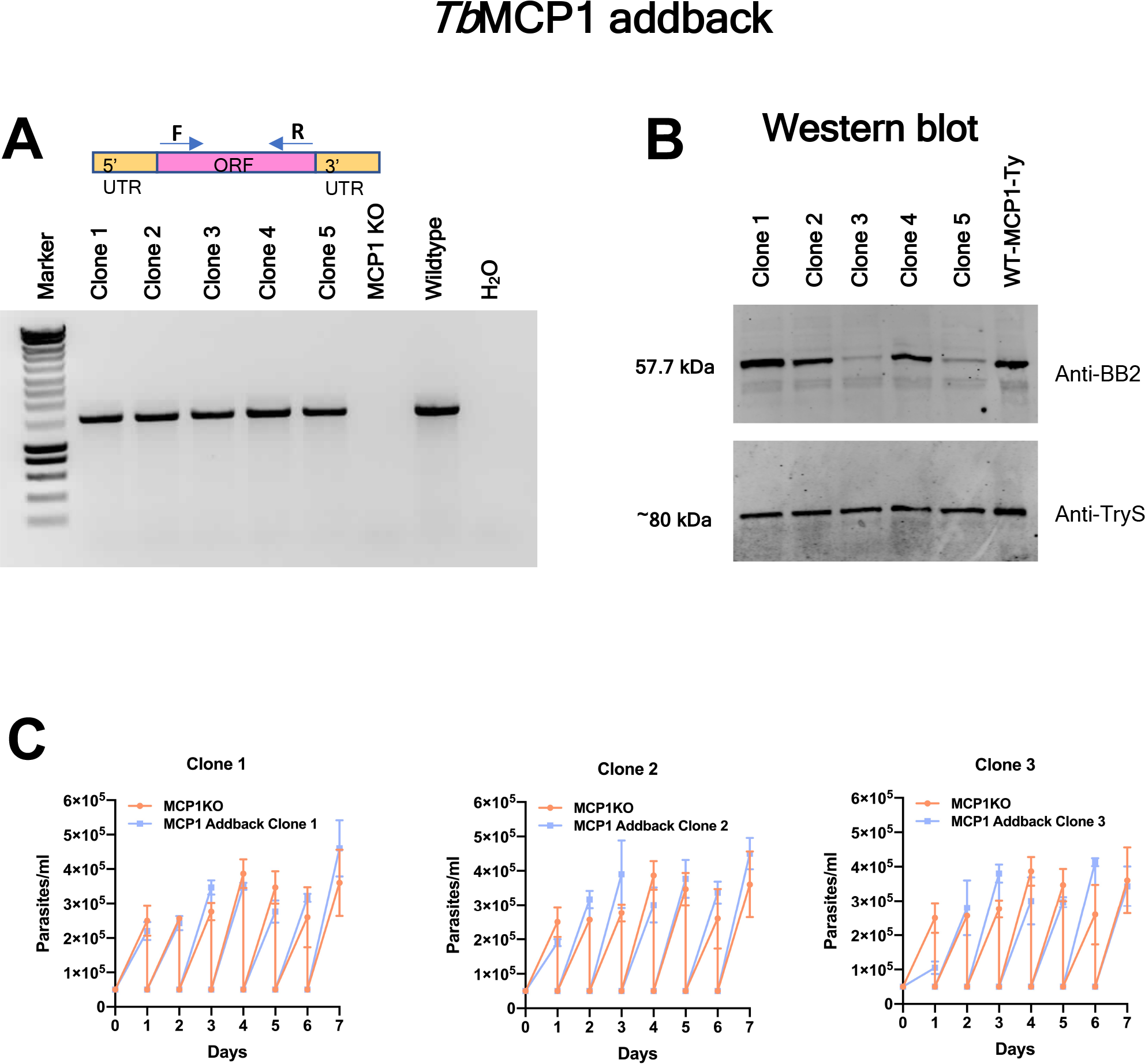
Validation of MCP1 add-back exprression and growth profile in vitro. A. PCR analysis of Tb927.11.2500 null mutants with a restored, add back, gene copy. Gene specific primers amplify the Tb927.11.2500 gene in the add back lines and parental cells, but not in null mutant. B. Western blot analysis confirming expression of the add back copy of Tb927.11.2500. The control line is parental parasites with an endogenously Ty1 epitope tagged copy of Tb927.11.2500 (WT-MCP1-ty). Equivalent loading is shown by trypanothione synthetase (TryS). C. *In vitro* growth profile of three add back clones in comparison to the null mutant. In each the growth *in vitro* of the cell lines was equivalent.

**Supplementary Figure 6.**
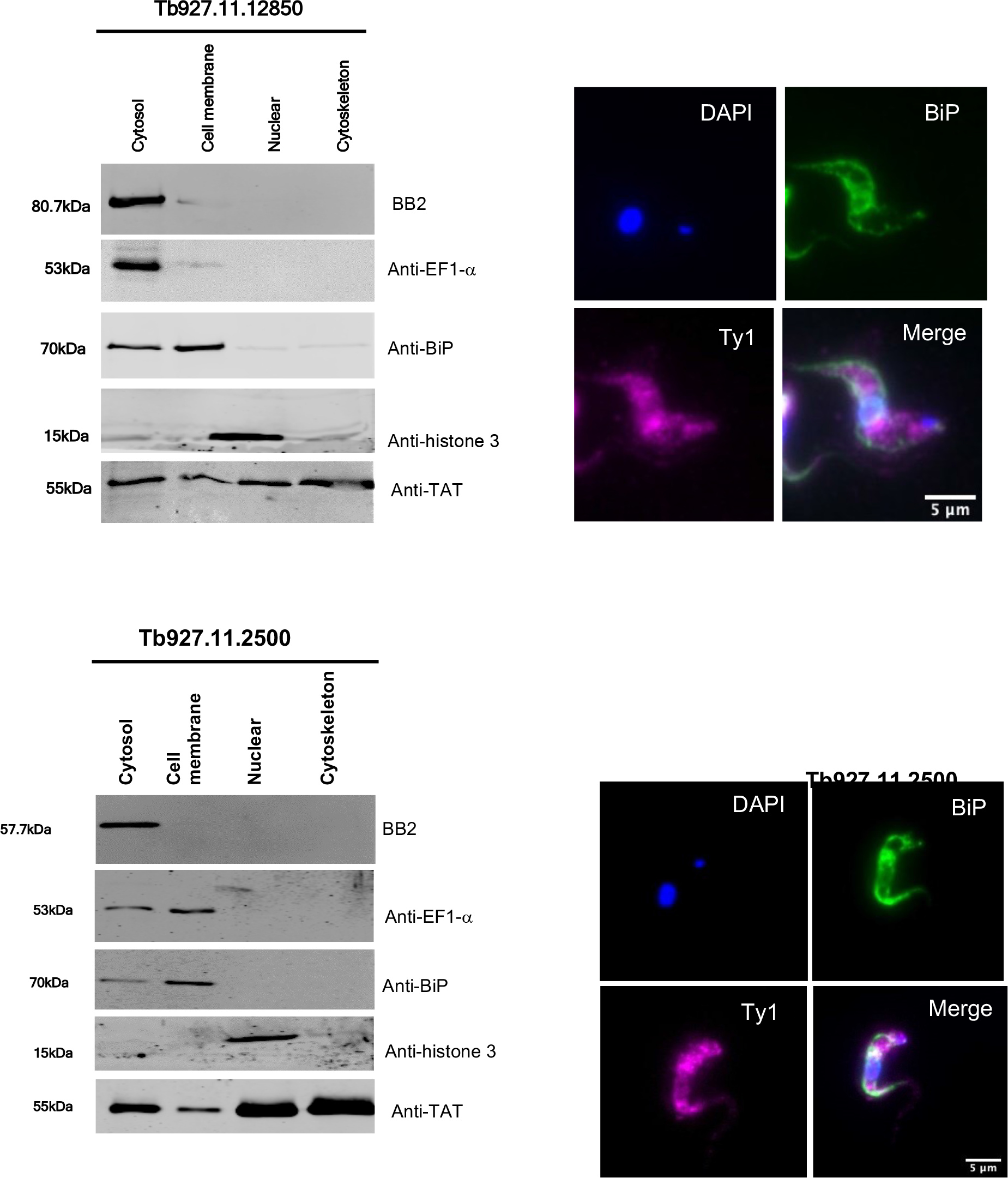
Oligopeptidase B and metallocarboxypeptidase 1 are cytosolic proteins. A. Cell fractionation of parasites expressing endogenously Ty1 epitope tagged Tb927.11.12850 and Tb927.11.2500. The tagged proteins (‘BB2’) were mostly cytosolically associated. Marker proteins were EF1-alpha (cytosol and some membrane fraction), BiP (some cytosolic and mainly ER), histone H3 (nuclear) and alpha tubulin (detected in all fractions). B. Immunofluorescence analysis of the distribution of endogenously Ty1 epitope tagged Tb927.11.12850 or Tb927.11.2500 with respect to the ER marker BiP. While BiP exhibited the expected reticulated pattern, the peptidases were uniformly distributed throughout the cell, reflective of cytosolic location.

